# Automated segmentation of skin strata in reflectance confocal microscopy depth stacks

**DOI:** 10.1101/022137

**Authors:** Samuel C. Hames, Marco Ardigò, H. Peter Soyer, Andrew P. Bradley, Tarl W. Prow

## Abstract

Reflectance confocal microscopy (RCM) is a powerful tool for in-vivo examination of a variety of skin diseases. However, current use of RCM depends on qualitative examination by a human expert to look for specific features in the different strata of the skin. Developing approaches to quantify features in RCM imagery requires an automated understanding of what anatomical strata is present in a given en-face section. This work presents an automated approach using a bag of features approach to represent en-face sections and a logistic regression classifier to classify sections into one of four classes (stratum corneum, viable epidermis, dermal-epidermal junction and papillary dermis). This approach was developed and tested using a dataset of 308 depth stacks from 54 volunteers in two age groups (20-30 and 50-70 years of age). The classification accuracy on the test set was 85.6%. The mean absolute error in determining the interface depth for each of the stratum corneum/viable epidermis, viable epidermis/dermal-epidermal junction and dermal-epidermal junction/papillary dermis interfaces were 3.1 *μm*, 6.0 *μm* and 5.5 *μm* respectively. The probabilities predicted by the classifier in the test set showed that the classifier learned an effective model of the anatomy of human skin.

## Introduction

Reflectance confocal microscopy (RCM) is capable of imaging human skin in-vivo at high resolution, with available machines capable of lateral resolutions of 0.5-1 *μm* [1]. RCM has been established as an effective tool for many applications in the assessment of human skin, such as the diagnosis of melanoma and keratinocyte skin cancers [2,3] and the assessment of inflammatory skin diseases [4]. However, appropriate training and experience are required to interpret RCM imagery: unlike the transverse sections of histopathology RCM images are acquired en-face and instead of the contrast provided by hematoxylin and eosin staining only a monochrome image showing variation in reflectance at one wavelength is available. While the non-invasive character of RCM imaging allows for the collection of large numbers of images, extracting quantitative information requires the time and expense of evaluation by a human expert. Fully exploiting the potential of RCM as a non-invasive imaging source requires new tools for standardised and streamlined assessment of large datasets.

While several approaches to standardised assessment using image analysis have been proposed none are currently in clinical use. These assessment approaches have focused on one of three main categories: 1) Quantifying and detecting specific features such as counting keratinocytes [5], detecting pagetoid cells [6] and evaluating photoageing [7], 2) Computer aided diagnosis of malignant melanocytic lesions [8] and 3) Identifying the anatomical structures of human skin [9,10]. Although both Somoza et al. and Kurugol et al. considered the problem of understanding human skin their work is limited: Kurugol et al. consider only the location of the dermal-epidermal junction and showed good performance only in darker skin types. While Somoza et al. showed an approach for complete segmentation of human skin their results were only a pilot study on 3 stacks. To date there is no robust automated method for understanding the complete anatomical structure of human skin in RCM depth stacks.

Understanding the anatomical structure of the skin is one of the fundamental tasks for assessment of RCM and conventional histopathology. It is necessary firstly for examining gross changes, such as thickening of the viable epidermis or stratum corneum. Secondly, an understanding of the distinct strata is also needed to guide the search for specific features. For example examining the honeycomb pattern of keratinocytes in the viable epidermis as a feature of photoageing [11] requires knowing at what depth the viable epidermis occurs in a stack. Therefore automated segmentation of the anatomical strata of the skin is a valuable starting point for standardised and automated analysis of RCM depth stacks.

The aim of this study was to develop a tool for automatically segmenting the distinct anatomical strata of human skin. This paper will outline an approach for achieving this aim using a bag of features approach to representing en-face optical sections that uses a dictionary of visual features learned by clustering image patches extracted from en-face RCM sections. This approach was trained and validated by comparison with an expert human observer. A bag of features approach was selected because these models are capable of modelling textural features and make minimal assumptions about the nature of the problem in addition the representative features are learned directly from the image data and do not require manual selection of relevant features. Bag of features also have demonstrated success in a number of medical imaging applications including, for example, colorectal tumor classification in endoscopic images [12], X-ray categorization [13] and histopathology image classification [14].

## Methods

### Participants and data acquisition

This study was conducted according to the Declaration of Helsinki with approval from the Metro South Human Research Ethics Committee and written informed consent obtained from participants. Participants were recruited in two age groups to represent a spectrum of normal skin. A total of 57 participants were recruited: 25 aged 20-29 (14 female) and 32 aged 50-59 (18 female). For the 20-29 year old group there were 7 phototype I, 9 phototype II, 7 phototype III, 1 phototype IV and 1 with no data available. For the 50-59 year old group there were 9 phototype I, 9 phototype II, 12 phototype III, 1 phototype IV and 1 with no data available. Recruitment focused on phototypes I-III to study photoageing in the context of light skin types. This dataset thus forms an ideal testbed for algorithm development in the context of the most challenging phototypes to examine with RCM. This dataset forms a superset of the data from Australian participants examined for the signs of photoageing in [11] and also used in [7].

Dorsal and volar skin of the forearm was imaged using a Vivascope 1500 (Caliber I.D, Rochester, NY, USA). A depth stack of en-face optical sections with vertical spacing of 2 *μm* was acquired to a depth of at least 50 sections (100 *μm*). Each en-face section had a field of 500 *μm* by 500 *μm* and 1000 by 1000 pixels. At least 2 stacks were acquired for each participant and body site, leading to a dataset of 335 stacks.

### Strata definition and labelling

For the purposes of segmentation four distinct anatomical strata were identified in the skin: the stratum corneum, viable epidermis, dermal-epidermal junction and the papillary dermis. To simplify the problem of establishing a ground truth in a large number of stacks it was assumed that a single anatomical strata was present in each en-face section and the extent of each strata was determined by identifying where three distinct features first occured within the stack, as illustrated in Fig. 1. This approach was inspired by work on weakly supervised video classification [15]. The specific features were based on [16]: visible honeycomb patterns corresponding to viable keratinocytes, visible bright bands corresponding to basal cells/dermal papillae and finally the absence of features of the basal layer -or- clearly visible fibrillar structures in the dermis. The en-face section with the first visible honeycomb pattern was considered the first layer of the viable epidermis - all sections above this layer were labelled as stratum corneum. En-face sections from the first honeycomb pattern to the first dermal papillae were labelled as viable epidermis. Sections from the first dermal papillae and up to, but not including, the en-face section where the features of the basal layer were no longer visible were labelled as dermal-epidermal junction. All sections below these were labelled as papillary dermis.

**Figure 1.**
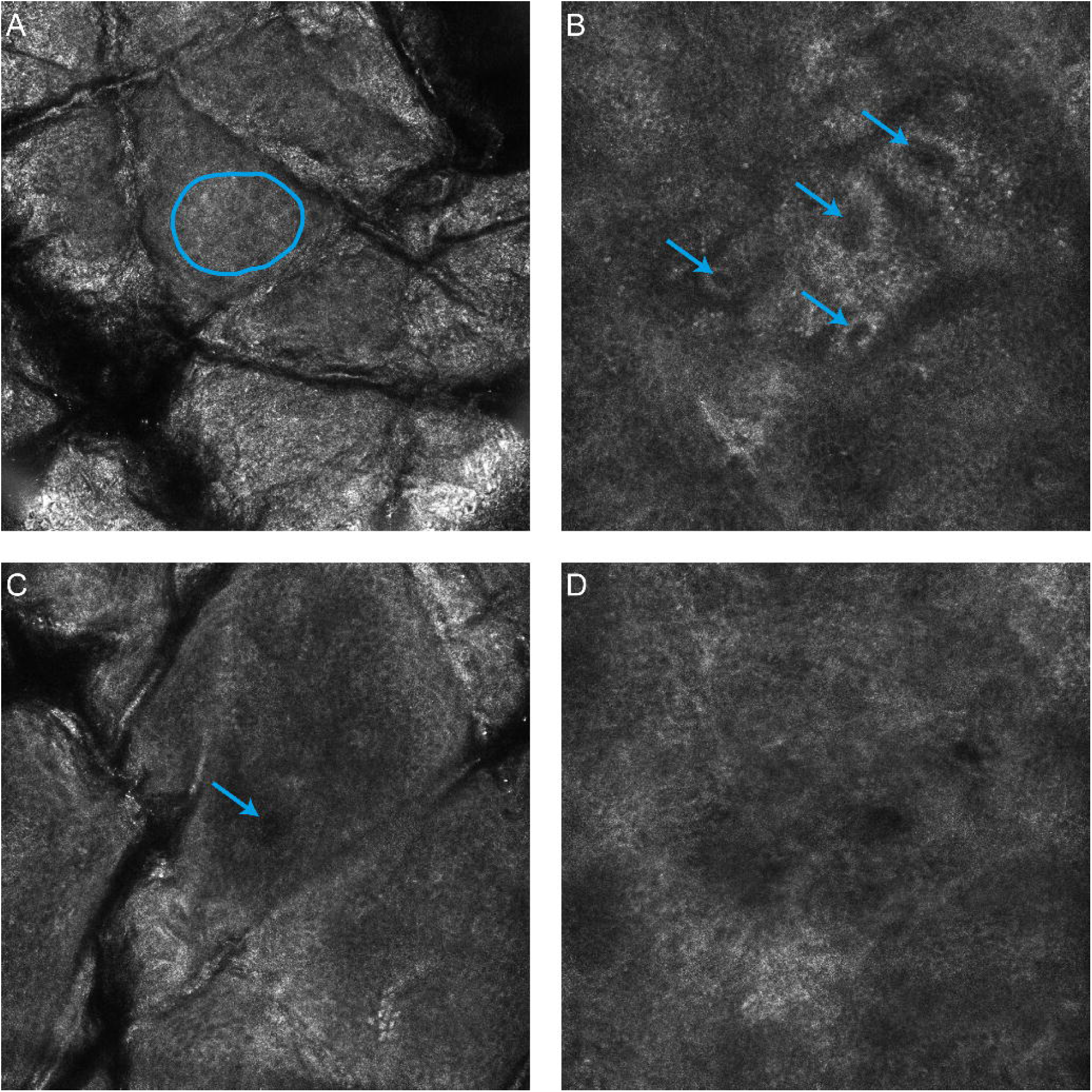
RCM features used in identification of the start and end of each anatomical strata. A) The start of the viable epidermis is marked by the first visible honeycomb pattern (Blue outline). B) The start of the dermal-epidermal junction is marked by the first clearly defined papillae with bright ring of basal cells surrounding a dark center (Blue arrows). C) The start of the papillary dermis is marked by the absence of features corresponding to the basal layer. D) Alternative definition of the start of the dermal-epidermal junction for light skin types, a dark featureless patch surrounded by keratinocytes with the absence of a bright basal layer (Blue arrow).

The dataset was labelled by a dermatologist with significant clinical experience with RCM (MA). Labelling was performed twice with four weeks in between to assess the intra-observer variability of the selected features. This second labelling was taken to be the ground truth for evaluation purposes due to the improved familiarity with the specific features and confidence in the results by the assessor. Stacks were organised by participant and bodysite and were labelled sequentially as would be performed in a clinical setting.

Some stacks showed motion or other artifacts making accurate labelling impossible - these stacks were excluded from the analysis. In addition if fewer than two examples for each body site were considered acceptable a participant was excluded from the analysis. A total of 16,144 images in 308 stacks from 54 participants could be successfully labelled and were included for further analysis.

### Representation of en-face sections using bags of features

A dictionary of representative visual features was learned using a process adapted from [17] using normalisation, whitening and spherical k-means to cluster small image patches drawn from the set of stacks. Initially the original high resolution optical sections were downsampled by a factor of 4 in each direction to 250 by 250 pixels (2 *μm* per pixel). This was necessary to match the scale of the extracted patches to the expected features in the image, and also to limit the dimensionality of features used in clustering. Following resizing a fixed number of patches were extracted from random locations in each en-face section. These patches were normalised to zero mean and unit variance (regularised by a constant to avoid singularities for low variance patches). These normalised patches were then whitened using a whitening matrix learned from the all extracted patches using the zero component analysis (ZCA) transform. Following [17] a regularisation factor, ε, was used, and additionally, only the top eigenvectors such that 95% of the energy was preserved in the transform were retained [18]. This last step was necessary to avoid numerical instability due to a number of low energy principle axes. After whitening the dictionary was learned by clustering using spherical k-means [19]. For encoding and clustering speed a hierarchical approach was applied to clustering, after [20,21]. A small fixed number of iterations (10) were used at each layer in the hierarchy, both for simplicity, and because of previous observations that few iterations are necessary for convergence [17].

Having learned a dictionary, each en-face section was then represented by the histogram of counts of visual features found in the image. Initially all raw patches in an en-face section were extracted in a dense sliding window fashion. After discarding zero variance patches, the extracted patches were normalised and whitened using the whitening matrix found while learning the dictionary. A whitened patch was quantized to the most similar (in a cosine sense) element in the dictionary. The final histogram was term-frequency normalised to be of unit length (L2 norm). The dataset was augmented (similar to [22]) by considering the original en-face section plus three rotations (at 90, 180 and 270 degrees). In the training phase these rotated examples plus the original section were considered as separate en-face sections while at testing phase the histograms from each rotation were pooled by averaging.

The histograms of visual feature counts for each en-face section were finally classified using L1 regularised logistic regression from the scikit-learn package [23]. The multiple classes were handled using a one-vs-rest scheme. Logistic regression was selected as a classifier because it can output meaningful probability estimates. L1 regularisation was selected for its demonstrated performance in handling cases with large number of irrelevant features [24], as might be expected to occur when learning a large number of features using a simple clustering methodology.

### Parameter estimation and performance assessment

A stratified random sample of participants from the two age groups (18 of the 54 participants, 8 from the 20-29 age group, 10 from the 50-59 age group) was held out as a final test set. The remainder were used for training and parameter estimation. Ten fold cross validation partitioned on the participants was then used on the training set with a coarse grid search to determine the parameters for maximum mean classifier accuracy over all of the folds. The grid search investigated the number of layers (1-3) and the number of splits at each layer (4-18) in the hierarchical dictionary as well as the logistic regression regularisation constant C (10^0^, 10^1^, … 10^5^). The total number of patches extracted for the dictionary learning process was constrained so that it could be performed within an 8GB memory limit. Other parameters were held constant: the size of extracted patches (7x7 pixels), the regularisation parameters for the patch normalisation (1) and the whitening transform (0.1), the energy retained in the ZCA transform (95%) and the number of iterations of the spherical k-means algorithm (10).

The classification performance of the algorithm was measured by applying the dictionary learning, encoding and classifier training process to the complete training set (with the optimal parameters selected by the grid search), then applying the learned feature encoding and classifier to the test set. The test set was not used at any point for training, validation or tuning. The output of the multi-class classifier was assessed by calculating the accuracy over all classes and the confusion matrix between the dermatologist labelling and the automatic classifications. The accuracy was also calculated for combinations of age and bodysite with strata type or phototype. To ensure that the classification was reliable for all stacks and not just on average the accuracy of the classifier was also calculated per stack in the test set.

Clinical utility was further assessed by using the classification output on independent sections to estimate the interface locations between each strata. Each interface location was determined by counting the number of sections of an anatomical strata and all strata that would occur above this. This number of layers is converted to a depth by multiplying by the depth between en-face sections. For example, the interface between the viable epidermis and the dermal-epidermal junction would be estimated as the sum of the number of sections classified as stratum corneum and viable epidermis multiplied by *μm*. Agreement between the dermatologist and automatically identified interfaces was measured by calculating the mean absolute error in interface position, as well as the Pearson correlation coefficient between the interface depths.

## Results

The classification accuracy over the entire test set was 85.6%. The parameters that optimised classification accuracy were a three layer encoding with 18 splits at each level (5832 total visual features) and C of 100. In comparison the intra-observer agreement of the dermatologist was 97.4%. The confusion matrices showing how images of each anatomical strata were classified by both the dermatologist and the automated method, are shown in Table 1. Table 2 shows the classification accuracies broken down by combinations of age, bodysite, strata and phototype. The complete classification results for the 5319 en-face sections in the test set are given in S1 Table.

**Table 1.** Comparison of human intraobserver agreement with automated approach on the 5319 sections in the test set.

|  |  | Dermatologist (1st Label) |  |  |  | Automatic |  |  |  |
| --- | --- | --- | --- | --- | --- | --- | --- | --- | --- |
|  |  | SC | VE | DEJ | PD | SC | VE | DEJ | PD |
| **Dermatologist (2nd Label)** | **SC** | 1292 | 1 | 0 | 0 | 1209 | 58 | 3 | 23 |
|  | **VE** | 23 | 1285 | 9 | 0 | 71 | 1085 | 154 | 7 |
|  | **DEJ** | 0 | 56 | 1523 | 22 | 0 | 143 | 1298 | 160 |
|  | **PD** | 0 | 0 | 26 | 1082 | 3 | 0 | 143 | 962 |

Key: SC = Stratum Corneum, VE = Viable Epidermis, DEJ = Dermal-epidermal Junction, PD = Papillary Dermis.

**Table 2.** Percentage of correctly classified sections by the automated approach in each subgroup.

|  |  | Strata |  |  |  | Phototype |  |  |  |
| --- | --- | --- | --- | --- | --- | --- | --- | --- | --- |
|  |  | SC | VE | DEJ | PD | I | II | III | Other |
| **20-30 years** | **Dorsal** | 95.7 | 82.8 | 88.7 | 91.2 | 90.9 | 84.4 | 90.0 | - |
|  | **Volar** | 91.2 | 79.7 | 85.1 | 62.9 | 80.2 | 84.0 | 80.0 | - |
| **50-70 years** | **Dorsal** | 91.9 | 82.2 | 76.8 | 92.5 | 80.3 | 94.0 | 84.6 | 86.0 |
|  | **Volar** | 95.6 | 84.2 | 77.0 | 92.6 | 85.9 | 88.7 | 89.3 | 87.6 |
| **Phototype** | **I** | 88.8 | 85.2 | 79.2 | 88.4 |  |  |  |  |
|  | **II** | 95.2 | 80.6 | 85.9 | 87.4 |  |  |  |  |
|  | **III** | 95.7 | 83.0 | 80.6 | 82.3 |  |  |  |  |
|  | **Other** | 95.5 | 77.4 | 81.0 | 92.9 |  |  |  |  |

Key: SC = Stratum Corneum, VE = Viable Epidermis, DEJ = Dermal-epidermal Junction, PD = Papillary Dermis.

The classification accuracy for each stack, and the agreement between the automatic and dermatologist identified interfaces between strata are shown in Fig. 2. The average absolute error and standard deviation in locating all interfaces was 4.8 ± 4.8 *μm* The average absolute error and standard deviation of the error in locating each interface was 3.1 ± 3.3 *μm*, 6.0 ± 5.3 *μm* and 5.5 ± 5.0 *μm* for the interfaces between the stratum corneum/viable epidermis, viable epidermis/dermal-epidermal junction and dermal-epidermal junction/papillary dermis respectively. The complete classification results for each stack in the test set are given in S2 Table.

**Figure 2.**
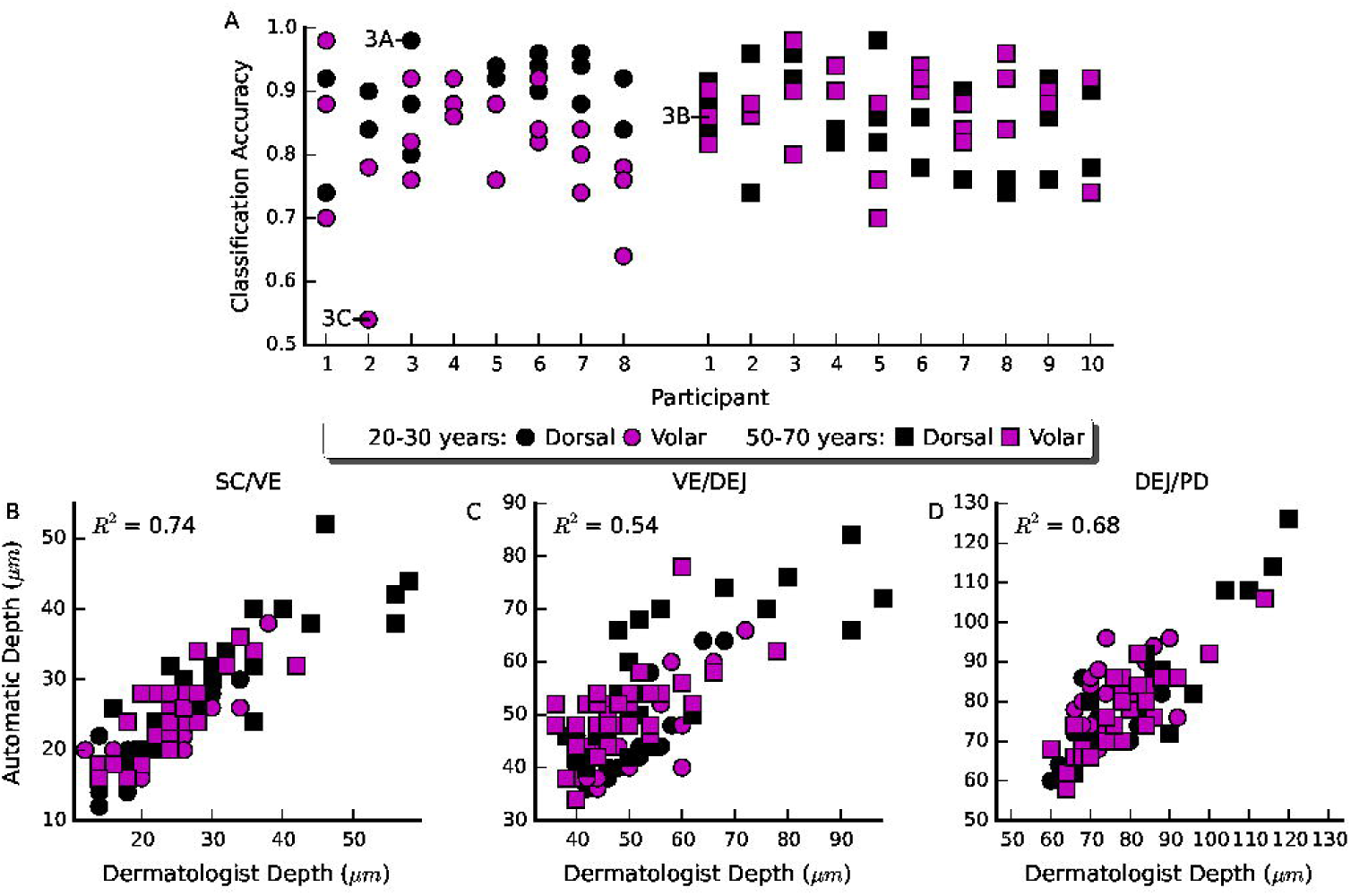
Accuracy and agreement of the automated approach with the dermatologist for individual stacks. A) Classification accuracy across all test stacks, organised by participant. The annotated examples are the best, median and worst accuracy stacks shown in Fig. 3. B-D) The correlation between the dermatologist identified interface and the automatically identified interface for each of the stratum corneum/viable epidermis, viable epidermis/dermal-epidermal junction and dermal-epidermal junction/papillary dermis interfaces.

Examples of automated and dermatologist identified interfaces are shown in Fig. 3. Also shown are the estimated strata probabilities through the depth of the stack, as estimated using the logistic regression classifier.

**Figure 3.**
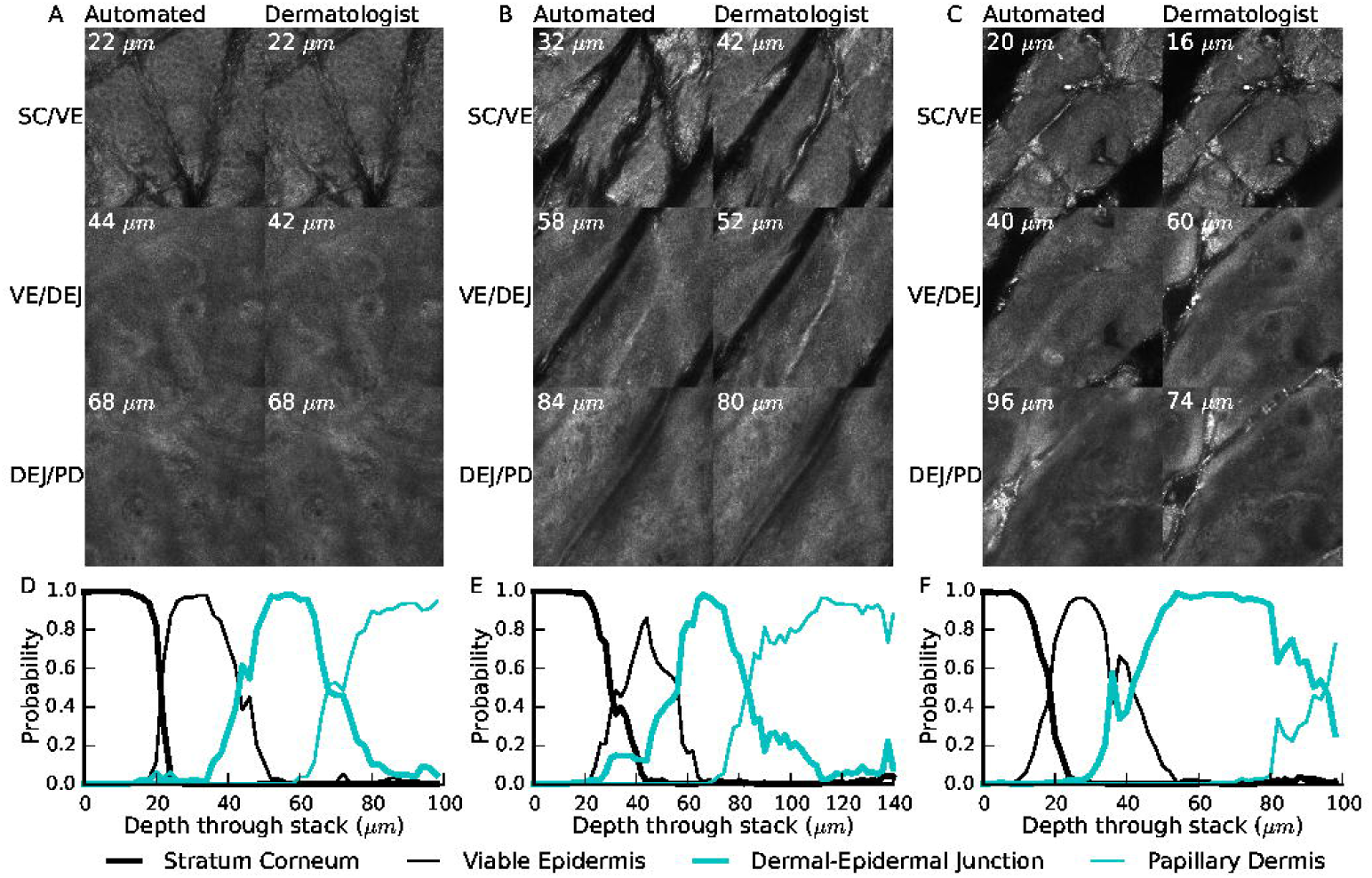
Example interfaces and probability output from the classifier. A-C) Comparison of the dermatologist identified interfaces with the automatically identified interfaces on the stacks with the highest, median and lowest accuracy respectively. D-F) The corresponding probability of each anatomical strata occurring in a section through the depth of the stack.

## Discussion

The visual features used to localize the anatomical strata within a stack were highly repeatable for the dermatologist observer (97.4% agreement between the first and second labelling sessions). In comparison the automated approach achieves a classification accuracy of 85.6% - a promising result for further automated analysis techniques. In particular the classification accuracy of the stratum corneum was excellent and did not strongly depend on the bodysite or age of the participant (Table 1.). In other strata performance was lower, but there did not appear to be a clear pattern in terms of bodysite or age. Similarly there did not appear to be a pattern for the classification accuracy of different phototypes. Since for all participants there is at least one stack classified with at least 90% accuracy (Fig. 2A), it is likely that the performance depends on some other characteristic of an individual stack. This other characteristic affecting performance could be the implicit assumption that a single strata is present in each en-face section: in the least accurately classified stack (Fig. 3C) there are furrows extending deep into the stack, causing a mixture of multiple anatomical strata present in each en-face section, whereas the most accurately classified example (Fig. 3A) exhibited only superficial furrows and approximately one anatomical strata per section. The stack with median accuracy (Fig. 3B) also exhibited deep furrows, but without bright reflectance and without strong mixing of strata.

Interpreting the results of the classification algorithm to identify the depths of different anatomical strata shows good agreement between the dermatologist and automated approach (Figure 2B-D). The mean absolute error in interface location was less than 6 *μm* for all three interfaces considered. In other words, on average less than three vertical en-face sections separated the dermatologist identified interface from the automatically identified interface. In the stratum corneum/viable epidermis in particular the average error was only 3.1 *μm* or just over 1.5 times the vertical depth spacing of en-face sections. Coupling this low absolute error with the high correlation between the dermatologist and automatically identified interface depth (*R*^2^ 0.74) suggests a possible use of this method to objectively quantify the thickness of the stratum corneum in-vivo.

Beyond the high classification accuracy the utility of this classification approach for further automated analyses is supported by examining the probabilistic output of the logistic regression classifier used (Fig. 3D-F). Despite being trained on individual sections the classifier effectively describes the order and transitions between different strata, with interface regions showing mixing of probabilities between different strata. Even in the least accurately classified stack, the estimated strata probabilities provide a useful approximate guideline as to where each anatomical strata is located within the stack.

Other work has analysed the structure of RCM depth stacks in different ways. Kurugol et al. [9] attempt to segment the full three dimensional shape of the dermal-epidermal junction based on low-level image features computed on image tiles. Unlike the approach presented here, where each en-face section is classified independently, they use smoothing in both the horizontal and vertical directions to ensure locally consistent results. Their algorithm was tested on 30 stacks from 30 participants, 15 light skin (phototype I-II) and 15 dark skin (phototype III-VI) - variations in skin appearance with aging and sun exposure were not considered. To analyse light phototypes they attempted to estimate a ‘transition’ region that approximately contained the dermal-epidermal junction - similar to the volume containing the dermal-epidermal junction reported here. Although the performance measures are not directly comparable because of the different classification methodologies, the average error in determining the location of the dermal-epidermal junction reported here (4.8 *μm*) is similar to their average error across all stacks in determining the location of the dermal-epidermal junction (8.5 *μm*).

Similar to this work Somoza et al. [10] show a method for classifying en-face sections as a single distinct strata. They define five strata from the stratum corneum to the papillary dermis. To classify en-face sections they use vector quantisation of histograms of visual features. The visual features were derived from a small set of hand selected filters, including oriented derivative of gaussians and laplacian of gaussians. In comparison, this work learns the features from the RCM sections themselves using unsupervised clustering. Although they show promising correlations between human and automated assessment (correlation coefficients 0.84-0.95), they examine only three RCM depth stacks and it is not clear if their work will generalise to other stacks. By comparison this work reports results on a much larger series of stacks and uses a robust validation approach with an independent test set used for final performance evaluation.

The method presented here is limited by two factors. Firstly, only dorsal and volar skin of the forearm is considered. This limitation is mitigated by the inclusion of multiple age-groups and a selection of phototypes, both factors that lead to observable changes in the skin. Secondly the focus is limited to phototypes I-III. However, this limitation is necessary to focus on the most challenging examples for human observers to examine, and it has already been shown that the improved reflectance of darker skin types are easier to examine automatically [9]. Since the method presented here incorporates brightness and contrast normalisation and focuses on learned features it is reasonable to expect that it will also work with phototypes IV and above.

In conclusion, it has been shown that a bag of features model is an effective tool for automatically segmenting and analysing reflectance confocal microscopy depth stacks. The classifier accuracy appears sufficient for approximate segmentation of RCM depth stacks across all four identified strata. Furthermore, in the stratum corneum it may be sufficiently accurate to allow automatic measurement of stratum corneum thickness with accuracy similar to a dermatologist. Future studies will focus on improving classification accuracy in strata other than the stratum corneum as well as assessment of skin pathologies including neoplastic and inflammatory skin diseases such as actinic keratosis and psoriasis.

## Supporting Information

S1 Table. Classification results for all 5319 sections in the test set.

S2 Table. The classification results and interface locations for each stack in the test set.

